# Strain- and age-dependent features of the nigro-striatal circuit in three common laboratory mouse strains, C57BL/6J, A/J, and DBA/2J - *Implications for Parkinson’s disease modeling*

**DOI:** 10.1101/2020.11.30.404293

**Authors:** Mélanie H. Thomas, Mona Karout, Beatriz Pardo Rodriguez, Yujuan Gui, Christian Jaeger, Alessandro Michelucci, Heike Kollmus, Klaus Schughart, Djalil Coowar, Rudi Balling, Michel Mittelbronn, Lasse Sinkkonen, Pierre Garcia, Manuel Buttini

## Abstract

Mouse models have been instrumental in understanding genetic determinants of aging and its crucial role in neurodegenerative diseases. However, few studies have analyzed the evolution of the mouse brain over time at baseline. Furthermore, mouse brain studies are commonly conducted on the C57BL/6 strain, limiting the analysis to a specific genetic background. In Parkinson’s disease, the gradual demise of nigral dopaminergic neurons mainly contributes to the motor symptoms. Interestingly, a decline of the dopaminergic neuron function and integrity is also a characteristic of physiological aging in some species. Age-related nigro-striatal features have never been studied in mice of different genetic backgrounds. In this study, we analyze the morphological features in the striatum of three common mouse strains, C57BL/6J, A/J, and DBA/2J at 3-, 9- and 15 months of age. By measuring dopaminergic markers, we uncover age-related changes that differ between strains and evolve dynamically over time. Overall, our results highlight the importance of considering background strain and age when studying the murine nigro-striatal circuit in health and disease.

**Highlights:**

- Study of the integrity of the nigro-striatal circuit in C57BL/6J, A/J, and DBA/2J at different ages
- Age related evolution of essential features of nigral dopaminergic neurons differ between strains
- Consider background strain and age is crutial to study the nigrostriatal circuit in health and disease

## 1. Introduction

Inbred mouse strains, as a consequence of at least 20 generations of brother-sister mating, are made of individual animals that are, for the most part, genetically identical and homozygous at all loci (Guenet and Benavides, 2011). This homogeneity makes them an essential tool in many facets of neuroscience research, from the study of basic central nervous system (CNS) functions over CNS development to, importantly, CNS diseases (Ellenbroek and Youn, 2016; van der Staay et al., 2009). The use of different mouse strains enables the study of how strain-specific genetic backgrounds influence CNS features and susceptibility to pathologies (McLin et al., 2006; Mulligan et al., 2018; Torres-Rojas et al., 2020). Because CNS disease manifestations and outcomes vary greatly between human individuals, such studies hold translational potential to help uncover mechanisms of neurodegeneration as well as neuroprotection. Various inbred strains have been used to study susceptibility to excitotoxicity (McLin et al., 2006; Schauwecker et al., 2004), seizures (Mohajeri et al., 2004), addiction and substance abuse (Korostynski et al., 2006; Mulligan et al., 2018), experimental autoimmune encephalomyelitis (McCarthy et al., 2012), and neurotoxin-induced Parkinson’s disease (PD) (Sedelis et al., 2000; Torres-Rojas et al., 2020).

PD is a chronic neurological illness with progressively worsening motor and non-motor symptoms (Poewe et al., 2017). Motor symptoms are caused by the gradual loss of dopaminergic neurons in the substantia nigra pars compacta (SN) and their axonal projections, leading to dopamine deficiency in the striatum (Raza et al., 2019). The age of onset, severity and progression of disease, as well as response to therapy, differ among patients and are modulated by genetic risk factors, aging, and environmental exposures (Jankovic et al., 1990; Kaplan et al., 2014).

Evidence suggests that some of these differences can be modeled in mice. Indeed, several studies have pointed out strain differences in susceptibility to PD neurotoxins, such as 1-methyl-4-phenyl-1,2,3,6-tetrahydropyridine (MPTP) (Hamre et al., 1999), or paraquat (Yin et al., 2011), with some strains, such as C57BL/6J being highly susceptible, whereas others, such as BALB/c and DBA/2J more resistant. Strains of the BXD mouse reference population, a population derived from founder strains C57BL/6J and DBA/2J, show a wide variety of quantitatively different responses to different paraquat dosing protocols (Torres-Rojas et al., 2020). Interestingly, A/J mice, at baseline, show lower performance than C57BL/6J mice in behavioral tests measuring locomotor activity (Ingram et al., 1981; Kollmus et al., 2020), tests which are often used to measure the behavioral consequences of nigro-striatal brain lesions (Grealish et al., 2010; Mokry, 1995).

In addition to genetic factors, age is a major risk factor for PD (Reeve et al., 2014). There are many factors that could contribute to the increased vulnerability of nigral dopaminergic neurons with age, such as increased oxidative stress, mitochondrial dysfunction, alpha-synuclein accumulation, or decreased protein degradation (Reeve et al., 2014). Importantly, even during normal aging, there is evidence for a decline of nigral dopaminergic neuronal function and integrity. Thus, in otherwise neurologically healthy people, longitudinal imaging studies have revealed a gradual age-related loss of striatal dopamine transporter and dopamine receptors (Karrer et al., 2017; Volkow et al., 1996a), and post-mortem pathology studies showed a gradual loss of dopamine neurons (Ma et al., 1999; Rudow et al., 2008). Studies in aging non-human primates show that markers of nigral dopaminergic dysfunction and degeneration, known to occur in PD, markedly increase over age, possibly reducing the threshold to disease susceptibility (Collier et al., 2017). There are, however, few studies though looking at the effect of age on the nigro-striatal circuit in different mouse strains. The vast majority of neurotoxin studies use mice that are anywhere between 1- and 3-4 month-old. A study investigating the toxicity of MPTP on C57BL/6 (substrain not specified) and BALB/c mice (Filipov et al., 2009) confirmed, on the one hand, that only C57BL/6 mice are susceptible to this toxin at a young age. On the other hand, the same study showed that at old age, not only were the C57BL/6J mice more affected, but so were BALB/c mice, albeit lesser than C57BL/6 mice. Studies on mice that are transgenic or knockout for PD familial genes usually use cohorts of older mice (up to around 12 month-old or older), and some of these models show age-dependent degeneration of their nigro-striatal circuit (Magen and Chesselet, 2010). Genetically altered mice though are, for obvious practical reasons, typically composed of individuals of just one strain, which, in neuroscience, is most commonly C57BL/6. Because sporadic PD in humans occurs predominantly in aged individuals, but also varies greatly among those individuals, the paucity of studies looking at the nigro-striatal circuit features in aging mice with different genetic backgrounds is surprising.

Hence, we set out to study, at baseline, the integrity of the nigro-striatal circuit and its evolution over age, in three widely used inbred mouse strains: C57BL/6J, A/J, and DBA/2J. We aimed at establishing a stepping stone for more advanced disease susceptibility investigations, and enable researchers to take decisions on which strains and age to use for their investigations. We measured dopamine concentration as well as dopaminergic neuron-related proteins such as tyrosine hydroxylase (TH), dopamine transporter (DAT), vesicular monoamine transporter 2 (VMAT2) and aldehyde dehydrogenase 1 family member A1 (ALDH1A1) in the dorsal striatum of 3-, 9- and 15-months-old mice of all 3 strains. We uncovered dynamic strain- and age-related patterns of nigro-striatal circuit features. Our results point to intricate interactions of genetic background with age in shaping the features of the nigro-striatal circuit in mice.

## 2. Material and methods

### 2.1. Animals

In this study, we used three different inbred mouse strains, C57BL/6J, A/J, and DBA/2J, at the age of 3, 9 and 15 months, with 4 to 15 mice per group, with similar numbers of males and females. Aged mice groups were typically lower in number as few mice were kept alive until that age, and there was attrition. The mouse breeders were purchased from Charles River, France (the purveyor of Jackson mice in Europe) for C57BL/6J and DBA/2J mice, or from Jackson Laboratory for A/J mice. The breeding of study groups was performed in house, at the Animal Facility of University of Luxembourg (Esch-sur-Alzette, Luxembourg), or at the Helmholtz Centre for Infection Research (Braunschweig, Germany). The mice used in this study were within four generations of breeding. Our protocol was reviewed and approved by the Animal Experimentation Ethics Committee (AEEC), the ‘Niedersächsisches Landesamt für Verbraucherschutz und Lebensmittelsicherheit, Oldenburg, Germany’ (Permit Number: 33.19-42502-05-19A394: *Organ-und Blutentnahme von genetisch unterschiedlichen Mausstämmen zur Identifizierung von Genen, die zur Parkinsonerkrankung führen*) as the appropriate government agency. Our experiments followed the 3R’s requirements for Animal Welfare, following the European Communities Council Directive 2010/63/EU, the national guidelines of the animal welfare in Luxembourg (Règlement grand-ducal adopted on January 11^th^, 2013) and the national guidelines of the animal welfare law in Germany (BGBl. I S. 1206, 1313 and BGBl. I S. 1934). Animals were group-housed in a controlled environment (12 hours-light/dark cycle) and received food and water *ad libitum*.

After anesthesia of each mouse with an intraperitoneal injection of ketamine (150 mg/kg) and medetomidine (1mg/kg), intracardiac perfusion was performed with PBS (phosphate-buffered saline). The brain was extracted and one hemibrain was dissected into midbrain and striatum and immediately snap-frozen, while the other hemibrain was fixed in 4% paraformaldehyde (PFA) (Sigma-Aldrich, 16005-1KG-R) for 48h, then transferred to PBS with 0.5% sodium azide (as preservative) and stored at 4°C until cutting,

### 2.2. Dopamine measurement

The dopamine measurement was performed on 4 to 15 snap-frozen striata per group. Two different protocols were used to measure dopamine concentration in the striatum of each mouse, thus dopamine concentration is expressed as percentage of 3-month-old C57BL/6J mice, to compare outcomes of both protocols (Jaeger et al., 2015; Jager et al., 2016). Briefly, the striatum was first pulverized or directly homogenized with grinding beads before extracting the metabolites using a liquid-liquid extraction method with methanol, water and chloroform. After shaking and centrifugation steps, the phase containing the polar metabolites was evaporated. The derivatization of metabolites in dried samples was performed as described (see references above), and the samples were then analyzed by gas chromatography - mass spectrometry (GC-MS) with an Agilent 7890A or 7890B coupled to an Agilent 5975C or 5977A inert XL mass selective detector (Diegem, Belgium), depending on the method used. We used a multi-analyte detection using a quadrupole analyzer in selected ion monitoring mode for a sensitive and precise quantification of dopamine and the internal standard DA-d4 (2-(3,4-Dihydroxyphenyl)ethyl-1,1,2,2-d4-amine HCl; C/D/N Isotopes Inc.).

### 2.3. Immunofluorescence and quantitative image analysis

The hemibrains of 4 to 6 mice per group were fixed in 4% PFA for 48 h and stored at 4°C in PBS/0.2% sodium azide. Parasagittal free floating sections (50 μm) were generated using a vibratome (Leica; VT 1000S) and stored at −20°C in a cryoprotective medium (polyvinyl pyrrolidone 1% w/v in PBS/ethylene glycol 1:1). We selected 3 sections per mouse to perform immunofluorescence. ALDH1A1 was mainly present in the upper part of the dorsal striatum, and its expression differed from lateral to medial sections in C57BL/6J mice (Supplementary Figure S1A). Thus we selected intermediate sections, where expression is highest in C57BL/6J mice, for all the animals.

After washing in PBS with 0.1% Triton X-100 (PBST) and permeabilization with 1.5% Triton X-100 and 3% H202 in PBS, 3-4 sections/mouse were blocked in a solution of 5% bovine serum albumin (BSA) in PBST. The sections were then incubated overnight with rabbit anti-TH (1:1000, Millipore), rat anti-DAT (1:1000, Millipore, MAB369), rabbit anti-ALDH1A1 (1:500, Abcam, ab215996) and rabbit anti-VMAT2 (1:150, Abcam, ab70808) diluted in a solution of 2% BSA in PBST. After washing in PBST, the sections were incubated for 2 hours with Alexa fluor™ 488 goat anti-rabbit (1:1000, Invitrogen), Alexa fluor™ 647 anti-rat (1:1000, Invitrogen) or Alexa fluor™ 488 anti-rabbit mounted on slides and embedded in fluoromount (SouthernBiotech, AL).

Imaging was performed using a Zeiss AxioImager Z1 upright microscope, coupled to a “Colibri” LED system, and an Mrm3 digital camera for image capture using Zeiss ZEN 2 Blue software. All measurements were performed on blinded sections, and codes were broken only after measurements for all images for each of the markers were completed. Three images (223.8 x 167.7 μm each) of the dorsal striatal area on each section were taken at 40x magnification using the Apotome system to obtain optical planes without out of focus reflection. Image were captured at optical plane levels were antibody stainings were uniform. For each marker, acquisition parameters were kept identical for each of the collected pictures, and all samples were blinded. All pictures were converted into 8-bit Tiff files for image analysis, which was done with the IMAGE J public software (version 1.51j8) as described (Jensen, 2013). After manual thresholding to capture only immunopositive structures, in each individual image for each of the markers (TH; DAT; VMAT2; ALDH1A1), two measurements were taken: first, the % image area occupied by immunopositive signals, and second, the mean grey value, or mean pixel intensity (scale 0-255 for 8-bit TIFF images) for the totality of immunopositive signals (Buttini et al., 1999; Guirado et al., 2018). The “% image area occupied” gives a relative measure of the physically present structural component that contains the detected protein, in other words, the density of a specific antibody-positive structure (Buttini et al., 1999). For instance, measurement of TH-positive axons in the striatum yields a relative measure of the level of branches of TH-positive axons in that area (Masliah et al., 2000). The “mean intensity”, gives a relative measure of the amount of protein detected by the antibody staining (Buttini et al., 2005; Jensen, 2013). A stronger signal reflects more of the antibody-detected protein, and vice-versa. Both measures do not necessarily always go into the same direction.

The estimation of number of TH-positive neurons in the SN was done morphometrically on blinded sections, as described elsewhere (Ashrafi et al., 2017). This method has been validated by stereological estimates of TH-neuron number (see Suppl. Material in (Ashrafi et al., 2017)).

### 2.4. Statistical analysis

GraphPad Prism 8 software was used for the statistical analysis. The data (normally distributed, Shapiro-Wilk normality test) were analyzed by two-way ANOVA followed by post hoc Tukey’s multiple comparisons test. P values below 0.05 were considered significant.

We present the data as tables, embedded in the text, with “<” or “>” symbols indicating the qualitative differences together with their percentage difference and their p values, as well as regular scatter graphs, embedded in figures.

## 3. Results

Here, we present the results of different measurements that capture the evolution of the nigro-striatal circuit in inbred mouse strains as they age. Dopamine (DA) is the neurotransmitter released by the neurons of the SN that project mainly toward the dorsal striatum and control movement (Berke, 2018). TH is the enzyme that converts the amino acid tyrosine to L-DOPA (3.4-dihydroxyphenylalanine) in the dopamine synthesis cascade (Daubner et al., 2011). DAT drives the dopamine reuptake out of the synaptic cleft back into intracellular vesicles (Jaber et al., 1997). VMAT2 is another membrane transporter that captures monoamine neurotransmitters (dopamine, serotonin, norepinephrine, histamine) from the neuronal cytoplasm into intracellular vesicles (Eiden and Weihe, 2011). ALDH1A1 in the brain, where it is specifically expressed in midbrain dopaminergic neurons, is involved in the metabolism of dopamine and norepinephrine (Anderson et al., 2011).

Our striatal dopamine data for the 3-month-old C57BL/6J and A/J presented have already been shown in another context (Thomas, 2020). Those data being essential for the comparison with other age groups (9 and 15 months) and the DBA/2J, we include them again in the present study. Also, in the present paper, we measured striatal TH density on a separate set of sections of 3-month-old C57BL/6J and A/J mice, and confirmed our previous observation (Thomas, 2020) showing that A/J mice have lower striatal TH density than C57BL/6J at 3 months of age.

### 3.1. Strain- and age-dependent variations of dopamine concentration in the striatum

The statistical analysis of striatal dopamine concentration, measured by GC-MS (see Methods), revealed significant effects for both strain (F=9.649, p=0.0002) and age (F=6.911, p=0.0017) on that measure (Table 1, Figure 1).

**Figure 1.**
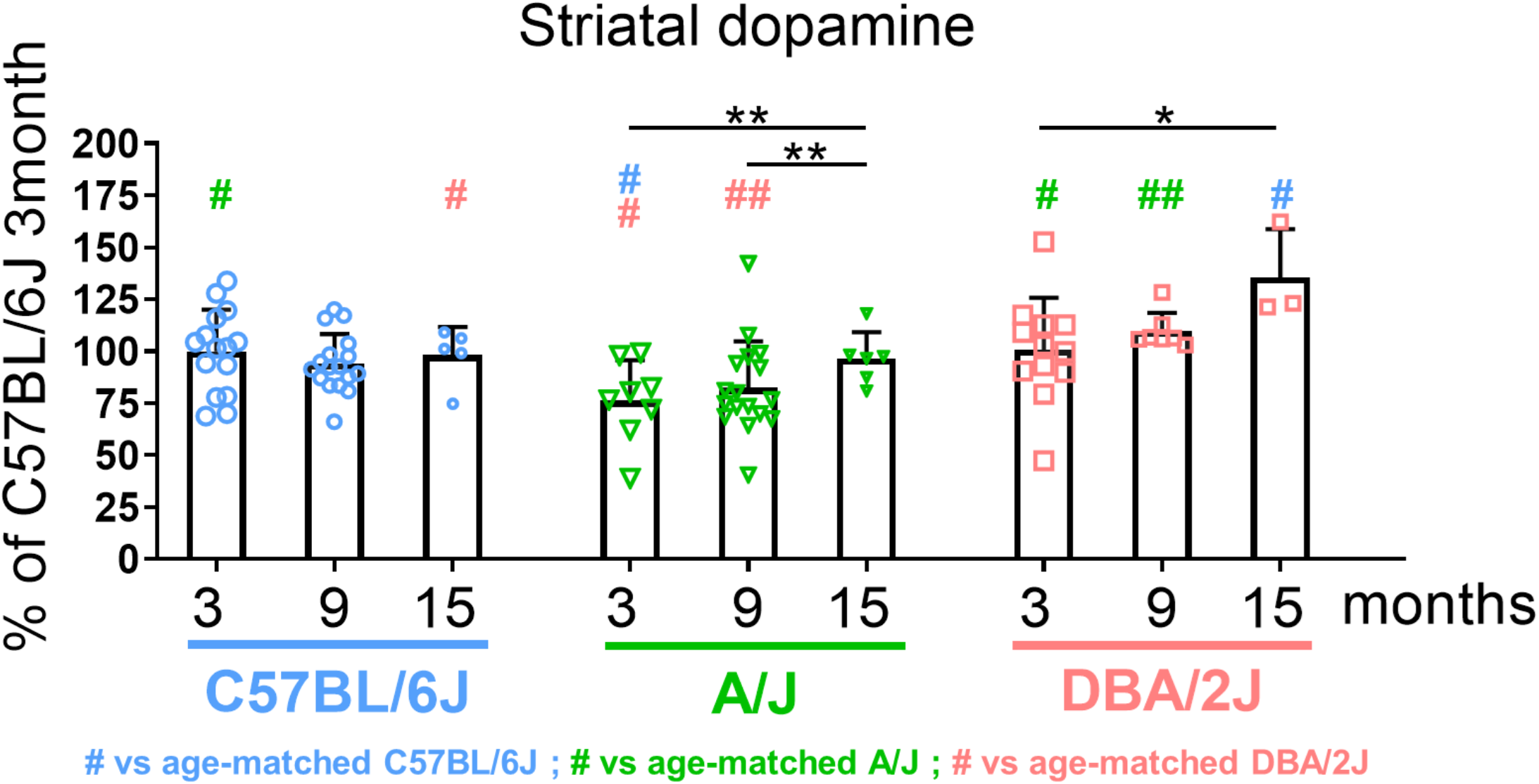
Striatal dopamine measured by GC-MS in 3-, 9- and 15-month-old C57BL/6J, A/J and DBA/2J mice. The dopamine level is shown as a percentage of C57BL/6J. Data are shown as means ± standard deviation, and analyzed with 2-way ANOVA followed by Tukey’s multiple comparisons test (*: p<0.05 and **: p<0.01 comparing the different age within each strain; #: p<0.05 and ##: p<0.01 comparing the different strains within each age, in blue for the comparisons to C57BL/6J, in green for the comparisons to A/J and in pink for the comparisons to DBA/2J).

**Table 1:** Level of dopamine in the dorsal striatum as a function of strain and age. BL6 stands for C57BL/6J, and DBA for DBA/2J. Tables lists comparison between strains at different ages (Strain), and between ages within each strain (Age).

|  |  | Striatal dopamine |  |  |
| --- | --- | --- | --- | --- |
|  |  | C57BL/6J vs A/J | C57BL/6J vs | A/J vs DBA/2J |
| Strain | 3 months | BL6 > A/J<br>23.8%, $p=0.0134$ | n.s. | A/J < DBA.<br>24.2%, $p=0.0152$ |
| | 9 months | n.s. | n.s. | A/J < DBA<br>28.1%, $p=0.005$ |
| | 15 months | n.s. | BL6 < DBA<br>27.6%, $p=0.0206$ | n.s. |
|  |  | 3 vs 9 months | 3 vs 15 months | 9 vs 15 months |
| Age | C57BL/6J | n.s. | n.s. | n.s. |
| | A/J | n.s. | 3M < 15M<br>21.1%, $p=0.0024$ | 9M < 15 M<br>18.3%, $p=0.005$ |
| | DBA/2J | n.s. | 3M < 15M<br>25.8%, $p=0.0137$ | n.s. |

The key observations were: 1. lower dopamine in the striatum of A/J mice at 3 and 9 months, but not 15 months, compared to both C57BL/6J and DBA/2J. 2. a significant uptick in dopamine in the striatum of both A/J and DBA/2J mice at 15 months compared to 3 or 9 months.

### 3.2. Strain and age-dependent variations of striatal TH-positive axons

The statistical analysis of TH-positive axons in the striatum, measured by quantitative image analysis on immunostained brain sections (see method), revealed significant main effects of both age on TH-positive axon density (F=12.54, p<0.0001) and strain (F=32.84, p<0.0001), but only of strain on TH-positive axon signal intensity (F=23.65, p<0.0001). There was also a significant interaction between strain and age on both measures (F=4.249, p=0.0056 for density, and F=7.327, p=0.0001 for intensity), indicating a differential effect of aging on TH axons between the strains. The results are shown in Table 2 and Figure 2.

**Figure 2.**
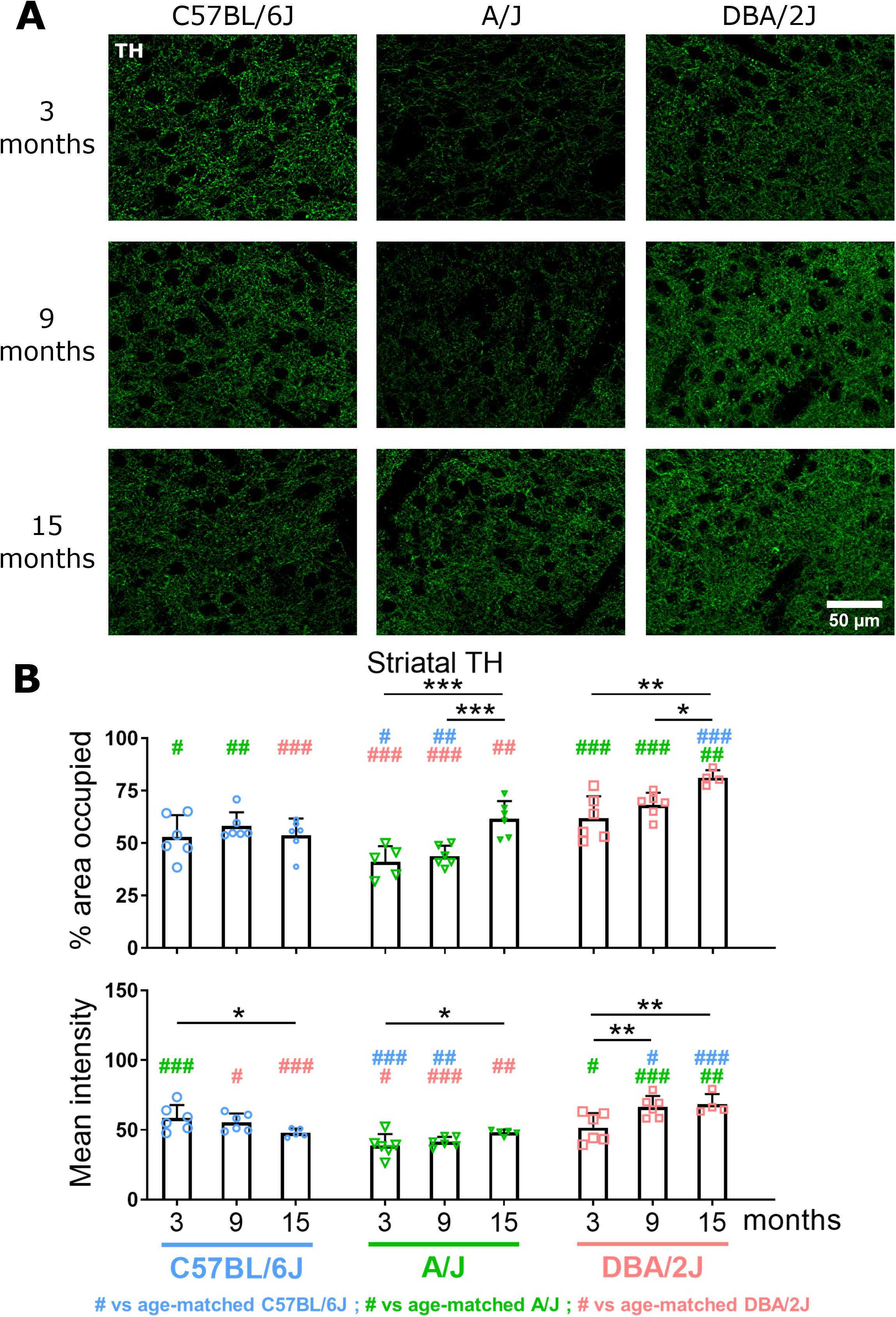
Striatal tyrosine hydroxylase (TH) measured by immunofluorescence in 3-, 9- and 15-month-old C57BL/6J, A/J and DBA/2J mice. (A) Images of TH staining, magnification 40X. (B) Quantification of % of area occupied (density) and intensity of TH. Data are shown as means ± standard deviation, and analyzed with 2-way ANOVA, followed by Tukey’s multiple comparisons test (*: p<0.05, **: p<0.01 and ***: p<0.001 comparing the different ages within each strain; #: p<0.05, ##: p<0.01 and ### < p<0.001 comparing the different strains within each age, in blue for the comparisons to C57BL/6J, in green for the comparisons to A/J and in pink for the comparisons to DBA/2J).

**Table 2:** Density and intensity of TH-positive striatal axons as a function of strain and age. BL6 stands for C57BL/6J, and DBA for DBA/2J. Tables lists comparison between strains at different ages (Strain), and between ages within each strain (Age).

|  |  | TH, % area (density) |  |  | TH, mean intensity |  |  |
| --- | --- | --- | --- | --- | --- | --- | --- |
|  |  | C57BL/6J vs A/J | C57BL/6J vs | A/J vs DBA/2J | C57BL/6J vs A/J | C57BL/6J vs | A/J vs DBA/2J |
| Strain | 3 months | BL6 > A/J<br>22.6%, $p=0.0382$ | n.s. | A/J < DBA.<br>14.1%, $p=0.0002$ | BL6 > A/J<br>34%, $p<0.0001$ | n.s. | A/J < DBA<br>25%, $p=0.009$ |
| | 9 months | BL6 > A/J<br>24.5%, $p=0.0078$ | n.s. | A/J < DBA<br>35.8%, $p<0.0001$ | BL6 > A/J<br>25.3%, $p=0.0044$ | BL6 < DBA<br>16.7%, $p=0.0282$ | A/J < DBA<br>37.8%, $p<0.0001$ |
| | 15 months | n.s. | BL6 < DBA.<br>33.9%, $p<0.0001$ | A/J < DBA.<br>24%, $p=0.0010$ | n.s. | BL6 < DBA.<br>29.9%, $p=0.0004$ | A/J < DBA<br>29.5%, $p=0.0004$ |
|  |  | 3 vs 9 months | 3 vs 15 months | 9 vs 15 months | 3 vs 9 months | 3 vs 15 months | 9 vs 15 months |
| Age | C57BL/6J | n.s. | n.s. | n.s. | n.s. | 3M > 15M<br>18%, $p=0.0497$ | n.s. |
| | A/J | n.s. | 3M < 15M<br>33.4%, $p=0.0002$ | 9M < 15M<br>28.9%, $p=0.0008$ | n.s. | n.s. | n.s. |
| | DBA/2J | n.s. | 3M < 15M<br>23.9%, $p=0.0011$ | 9M < 15M<br>15.9%, $p=0.0358$ | 3M < 9M<br>22.5%, $p=0.0024$ | 3M < 15M<br>24.5%, $p=0.0024$ | n.s. |

The key observations were: 1. In C57BL/6J mice, the TH-positive striatal axons are mostly stable with age, except for a loss of signal intensity at 15 months, indicating some age-related loss of TH. 2. In A/J mice, the TH-positive striatal axon density and intensity are lower on younger mice (3 and 9 months) than in their C57BL/6J and DBA/2J counterparts but are increased at 15 months. 3. In DBA/2J mice, both TH-positive striatal axon density and intensity increase substantially over age, reaching levels higher than in their C57BL/6J and A/J counterparts.

### 3.3. Strain and age-dependent variations of striatal DAT-positive synapses

The data analysis of DAT-positive synapses in the striatum (see method) showed significant effects of both strain and age on density of DAT-positive synapses (strain: F=12.87, p<0.0001; age: F=64.21, p<0.0001) as well as signal intensity of DAT (strain: F=13.16, p<0.0001; age: F=90, p<0.0001). There was also a significant interaction between strain and age (F=21.37, p<0.0001 for DAT density and F=15.27, p<0.0001 for DAT intensity), indicating a differential effect of aging on DAT-positive synapses between the strains. The results are shown in Table 3 and Figure 3.

**Figure 3.**
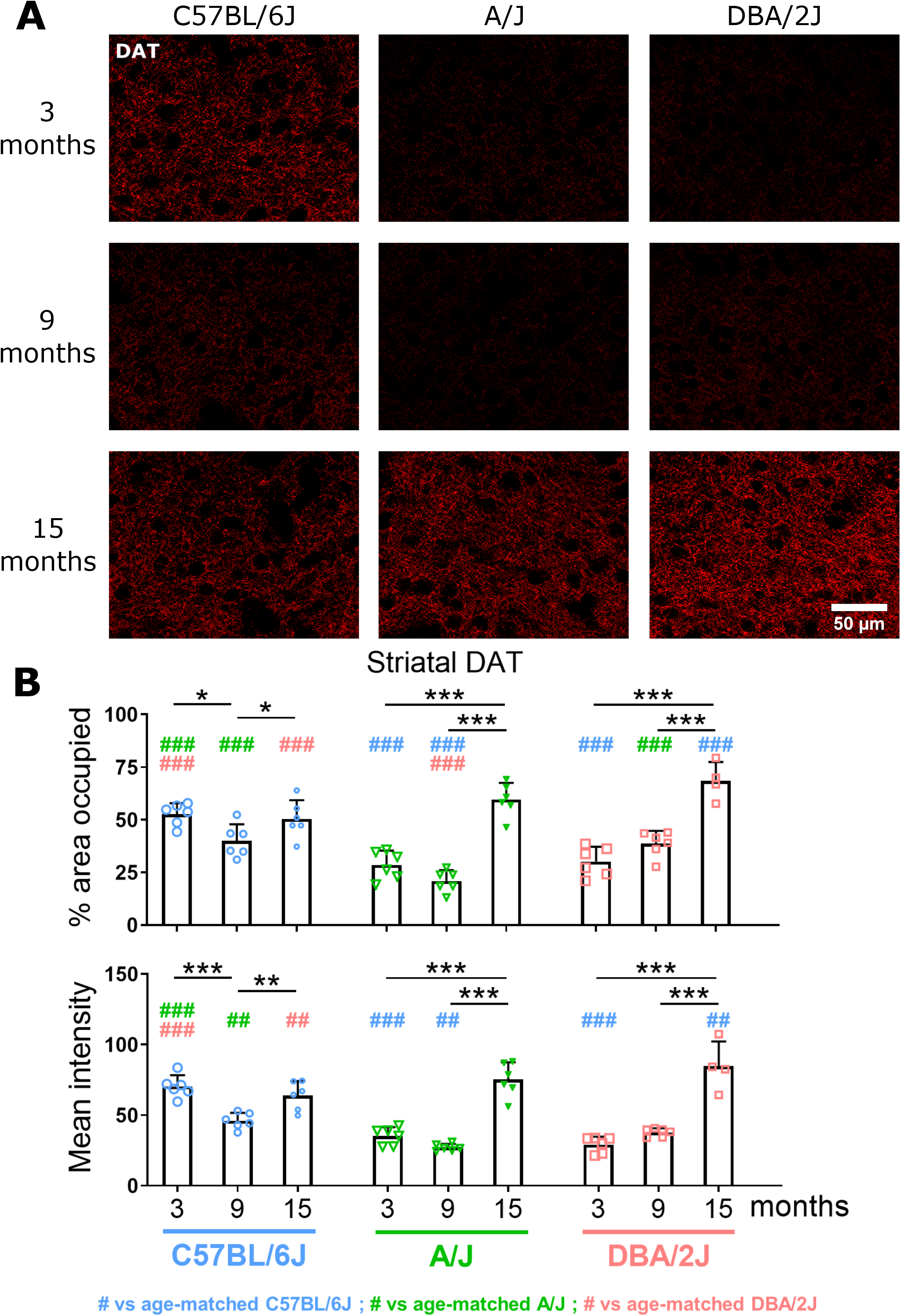
Striatal dopamine transporter (DAT) measured by immunofluorescence in 3-, 9- and 15-month-old C57BL/6J, A/J and DBA/2J mice. (A) Images of DAT staining, magnification 40X. (B) Quantification of % of area occupied and intensity of DAT. Data are shown as means ± standard deviation, and analyzed with 2-way ANOVA followed by Tukey’s multiple comparisons test (*: p<0.05, **: p<0.01 and ***: p<0.001 comparing the different ages within each strain; ##: p<0.01 and ### < p<0.001 comparing the different strains within each age, in blue for the comparisons to C57BL/6J, in green for the comparisons to A/J and in pink for the comparisons to DBA/2J).

**Table 3:** Density and intensity of DAT-positive striatal synapses as a function of strain and age. BL6 stands for C57BL/6J, and DBA for DBA/2J. Tables lists comparison between strains at different ages (Strain), and between ages within each strain (Age).

|  |  | DAT, % area (density) |  |  | DAT, mean intensity |  |  |
| --- | --- | --- | --- | --- | --- | --- | --- |
|  |  | C57BL/6J vs A/J | C57BL/6J vs | A/J vs DBA/2J | C57BL/6J vs A/J | C57BL/6J vs | A/J vs DBA/2J |
| Strain | 3 months | BL6 > A/J<br>45.7%, p<0.0001 | BL6 > DBA<br>42.7%, p<0.0001 | n.s. | BL6 > A/J<br>50%, p<0.0001 | BL6 > DBA<br>58.6%, p<0.0001 | n.s. |
|  | 9 months | BL6 > A/J<br>47.8%, p<0.0001 | n.s. | A/J < DBA<br>45.9%, p=0.0003 | BL6 > A/J<br>40.8%, p=0.0011 | n.s. | n.s. |
|  | 15 months | n.s. | BL6 < DBA<br>26.4%, p=0.0009 | n.s. | n.s. | BL6 < DBA<br>24.5%, p=0.0013 | n.s. |
|  |  | 3 vs 9 months | 3 vs 15 months | 9 vs 15 months | 3 vs 9 months | 3 vs 15 months | 9 vs 15 months |
| Age | C57BL/6J | 3M > 9M<br>23.9%, p=0.0104 | n.s. | 9M < 15M<br>20.4%, p=0.0423 | 3M > 9M<br>34.6%, p<0.0001 | n.s. | 9M < 15M<br>28%, p=0.0019 |
|  | A/J | n.s. | 3M < 15M<br>52%, p<0.0001 | 9M < 15M<br>64.8%, p<0.0001 | n.s. | 3M < 15M<br>53.3%, p<0.0001 | 9M < 15M<br>63.9%, p<0.0001 |
|  | DBA/2J | n.s. | 3M < 15M<br>59.9%, p<0.0001 | 9M < 15M<br>43.5%, p<0.0001 | n.s. | 3M < 15M<br>65.6%, p<0.0001 | 9M < 15M<br>55.3%, p<0.0001 |

The key observations were: 1. In 9-month-old C57BL/6J compared to their 3-month-old and 15-month-old counterparts, a transients lower striatal DAT density and intensity. 2. In younger (3 and 9 months) A/J and DBA/2J mice, significantly lower striatal DAT density and intensity than in their 15-month-old counterparts, and, in part, lower than their age-matched C57BL/6J counterparts. 3. The highest levels of striatal DAT density and intensity were found in 15-monthd-old DBA/2J, which is when they significantly surpassed those in age-matched C57BL/6J.

### 3.4. Strain and age-dependent variations of striatal VMAT2-positive structures

The statistical analysis of another transporter, VMAT2, in the striatum showed significant effects of strain and age on striatal VMAT2 density and intensity (F=11.48, p=0.0001, and F=3.751, p=0.0317 respectively), and only for strain (F=6.254, p=0.0042) for the VMAT2 density. There was also a significant interaction between strain and age (F=11.24, p<0.0001 for % of area and F=5.180, p=0.0017 for intensity), indicating a differential effect of aging on VMAT2-positive structures between the strains. The results are shown in Table 4 and Figure 4. The signals for VMAT2 were of two sorts: small, punctate and larger clusters of variable size (Figure 4, A). The first ones are most likely small synaptic vesicles, and the second ones intraneuronal dense core vesicles located in the trans-Golgi network (Nirenberg et al., 1995).

**Figure 4.**
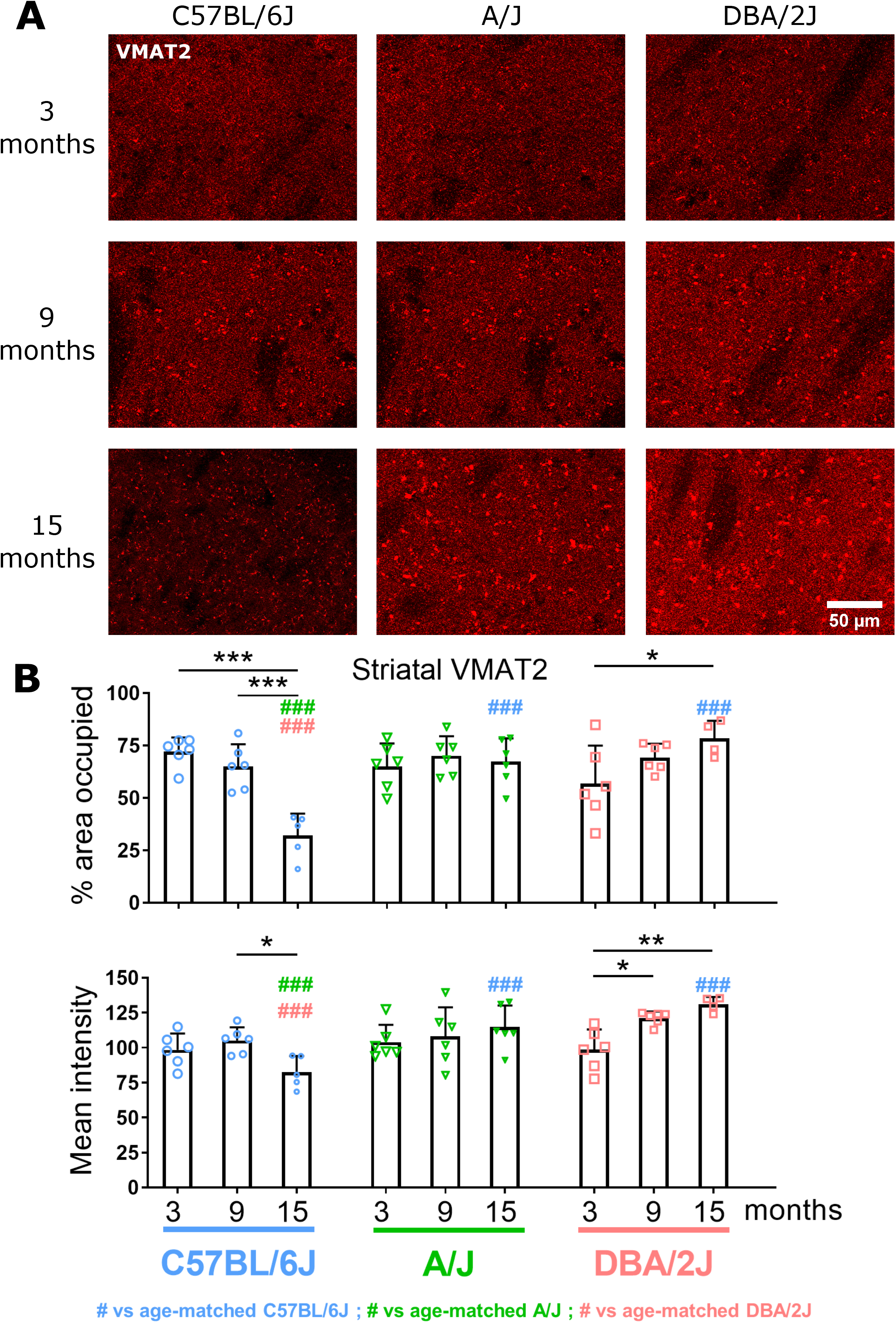
Striatal vesicular monoamine transporter 2 (VMAT2) measured by immunofluorescence in 3, 9- and 15-month-old C57BL/6J, A/J and DBA/2J mice. (A) Images of VMAT2 staining, magnification 40X. The larger clusters of signal are likely assemblies of large dense core vesicles at the trans Golgi network (Nirenberg et al.). (B) Quantification of % of area occupied (density) and intensity of DAT. Data are shown as means ± standard deviation, and analyzed with 2-way ANOVA followed by Tukey’s multiple comparisons test (*: p<0.05, **: p<0.01 and ***: p<0.001 comparing the different ages within each strain; ##: p<0.01 and ###: p<0.001 comparing the different strains within each age, in blue for the comparisons to C57BL/6J, in green for the comparisons to A/J and in pink for the comparisons to DBA/2J).

**Table 4:** Density and intensity of VMAT2-positive structures as a function of strain and age. BL6 stands for C57BL/6J, and DBA for DBA/2J. Tables lists comparison between strains at different ages (Strain), and between ages within each strain (Age).

|  |  | VMAT2, % area (density) |  |  | VMAT2, mean intensity |  |  |
| --- | --- | --- | --- | --- | --- | --- | --- |
|  |  | C57BL/6J vs A/J | C57BL/6J vs | A/J vs DBA/2J | C57BL/6J vs A/J | C57BL/6J vs | A/J vs DBA/2J |
| Strain | 3 months | n. s. | n. s. | n. s. | n. s. | n. s. | n. s. |
|  | 9 months | n. s. | n. s. | n. s. | n. s. | n. s. | n. s. |
|  | 15 months | BL6 < A/J<br>52.3%, p<0.0001 | BL6 < DBA<br>59.1%, p<0.0001 | n. s. | BL6 < A/J<br>28%, p=0.0005 | BL6 < DBA<br>36.8%, p<0.0001 | n. s. |
|  |  | 3 vs 9 months | 3 vs 15 months | 9 vs 15 months | 3 vs 9 months | 3 vs 15 months | 9 vs 15 months |
| Age | C57BL/6J | n. s. | 3M > 15M<br>55.5%, p<0.0001 | 9M > 15M<br>50.6%, p<0.0001 | n. s. | n. s. | 9M > 15M<br>21.6%, p=0.0164 |
|  | A/J | n. s. | n. s. | n. s. | n. s. | n. s. | n. s. |
|  | DBA/2J | n. s. | 3M < 15M<br>17.1%, p=0.0105 | n. s. | 3M < 9M<br>18.7%, p=0.0110 | 3M < 15M<br>24.8%, p=0.0010 | n. s. |

The key observation was: In the striatum of C57BL/6J, a significant drop of VMTA2 density and intensity at 15 months, whereas in that of A/J mice, both VMAT2 were stable, and in that of DBA/2J mice, they showed an age-dependent increase.

### 3.5. Strain and age-dependent variations of striatal ALDH1A1-positive structures

ALDH1A1 is expressed by a subset of neurons in the ventral tier of the SN, and the protein is primarily found in the rostral dorsal part of the striatum (Sgobio et al., 2017; Wu et al., 2019). This neuronal subpopulation particularly severely affected in PD (Cai et al., 2014). We confirmed the localization of striatal ALDH1A1 in the rostral dorsal part of the striatum (Supplementary Fig.S1), and quantified its relative level in this striatal subregion.

The data analysis of ALDH1A1-positive structures in the striatum showed significant main effects of both and strain (F=43.56, p<0.0001) age (F=32.76, p<0.0001) on ALDH1A1 density. There was also a significant interaction between strain and age for the ALDH1A1 density (F=35.03, p<0.0001) and intensity (F=31.88, p<0.0001), indicating a differential effect of aging on ALDH1A1-positive structures between the strains. The data are shown in Table 5 and Figure 5. Large-scale images of the striata of 3-, 9- and 15-months-old mice from the 3 strains are shown with a staining ALDH1A1 in Supplementary Figure S1B.

**Figure 5.**
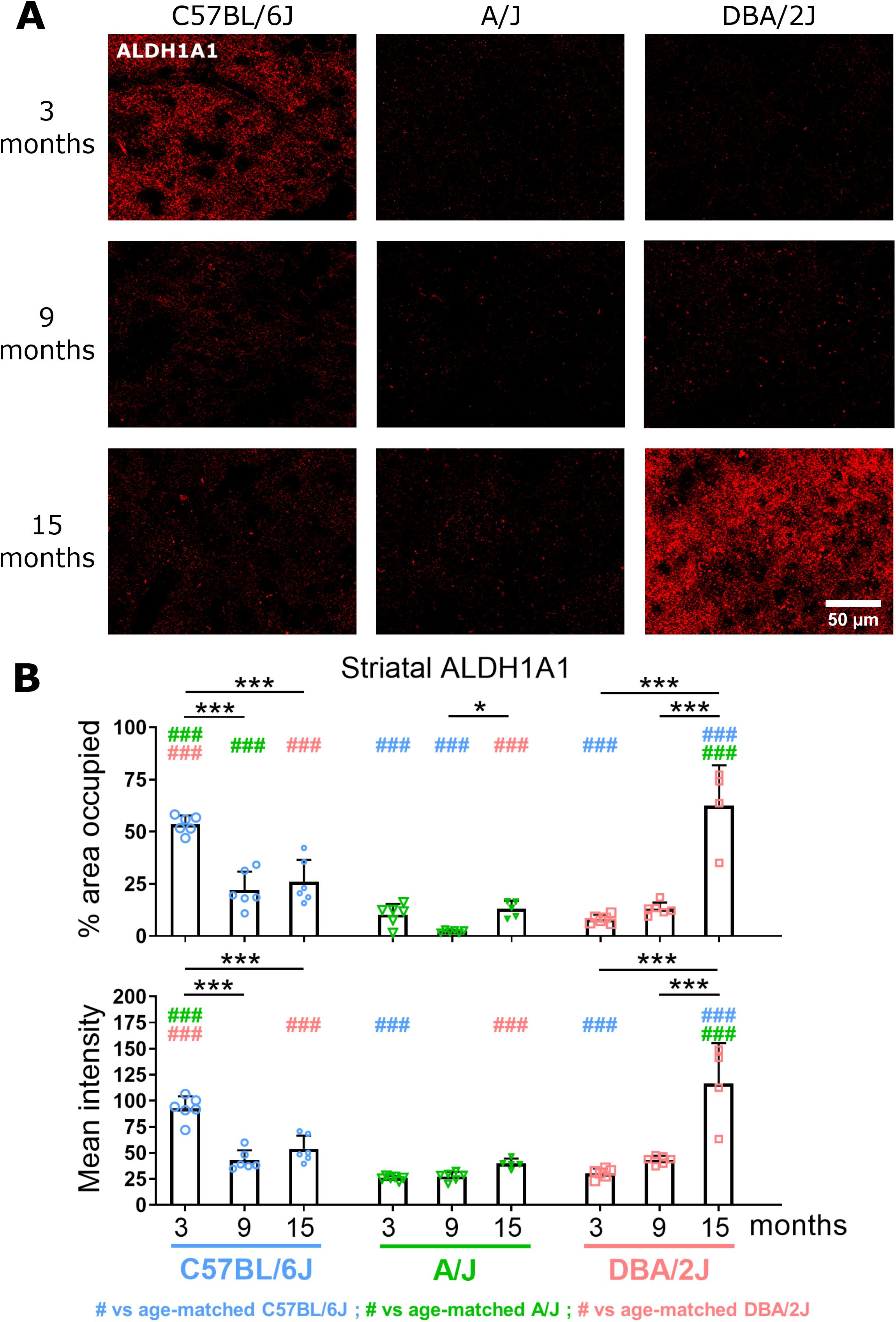
Striatal (rostral dorsal subregion) Aldehyde Dehydrogenase 1 Family Member A1 (ALDH1A1) measured by immunofluorescence in 3-, 9- and 15-month-old C57BL/6J, A/J and DBA/2J mice. (A) Images of ALDH1A1 staining, magnification 40X. (B) Quantification of % of area occupied and intensity of ALDH1A1. Data are shown as means ± standard deviation, and analyzed with 2-way ANOVA followed by Tukey’s multiple comparisons test (*: p<0.05, **: p<0.01 and ***: p<0.001 comparing the different ages within each strain; ##: p<0.01 and ###: p<0.001 comparing the different strains within each age, in blue for the comparisons to C57BL/6J, in green for the comparisons to A/J and in pink for the comparisons to DBA/2J).

**Table 5:** Density and intensity of ALDH1A1-positive structures as a function of strain and age. BL6 stands for C57BL/6J, and DBA for DBA/2J. Tables lists comparison between strains at different ages (Strain), and between ages within each strain (Age).

|  |  | ALDH1A1, % area (density) |  |  | ALDH1A1, mean intensity |  |  |
| --- | --- | --- | --- | --- | --- | --- | --- |
|  |  | C57BL/6J vs A/J | C57BL/6J vs | A/J vs DBA/2J | C57BL/6J vs A/J | C57BL/6J vs | A/J vs DBA/2J |
| Strain | 3 months | BL6 > A/J<br>80.7%, p<0.0001 | BL6 > DBA<br>84.8%, p<0.0001 | n.s. | BL6 > A/J<br>72.1%, p<0.0001 | BL6 > DBA<br>67.3%, p<0.0001 | n.s. |
|  | 9 months | BL6 > A/J<br>89.6%, p=0.0003 | n.s. | n.s. | n.s. | n.s. | n.s. |
|  | 15 months | n.s. | BL6 < DBA<br>58.4%, p<0.0001 | A/J < DBA<br>79.2%, p<0.0001 | n.s. | BL6 < DBA<br>53.9%, p<0.0001 | A/J < DBA<br>65.9%, p<0.0001 |
|  |  | 3 vs 9 months | 3 vs 15 months | 9 vs 15 months | 3 vs 9 months | 3 vs 15 months | 9 vs 15 months |
| Age | C57BL/6J | 3M > 9M<br>58.7%, p<0.0001 | 3M > 15M<br>51.4%, p<0.0001 | n.s. | 3M > 9M<br>53.8%, p<0.0001 | 3M > 15M<br>42%, p<0.0001 | n.s. |
|  | A/J | n.s. | n.s. | 9M < 15M<br>82.3%, p=0.0101 | n.s. | n.s. | n.s. |
|  | DBA/2J | n.s. | 3M < 15M<br>87%, p<0.0001 | 9M < 15M<br>79.2%, p<0.0001 | n.s. | 3M < 15M<br>74%, p<0.0001 | 9M < 15M<br>63%, p<0.0001 |

The key observations were: In the striatum of 3-month-old C57BL/6J and in that of 15-month-old DBA/2J, both ALDH1A1 density and intensity were higher than in all other groups, indicating an age-dependent decline of both measures in C57BL/6J mice but an increase in DBA/2J mice. In the striatum of A/J mice, ALDH1A1 remained stable.

### 3.6. Nigral dopaminergic neurons during aging in C57BL/6J, A/J, and DBA/2J strains

The number of TH-positive neurons in the SN was similar in all strains and stable over age (Figure 6).

**Figure 6.**
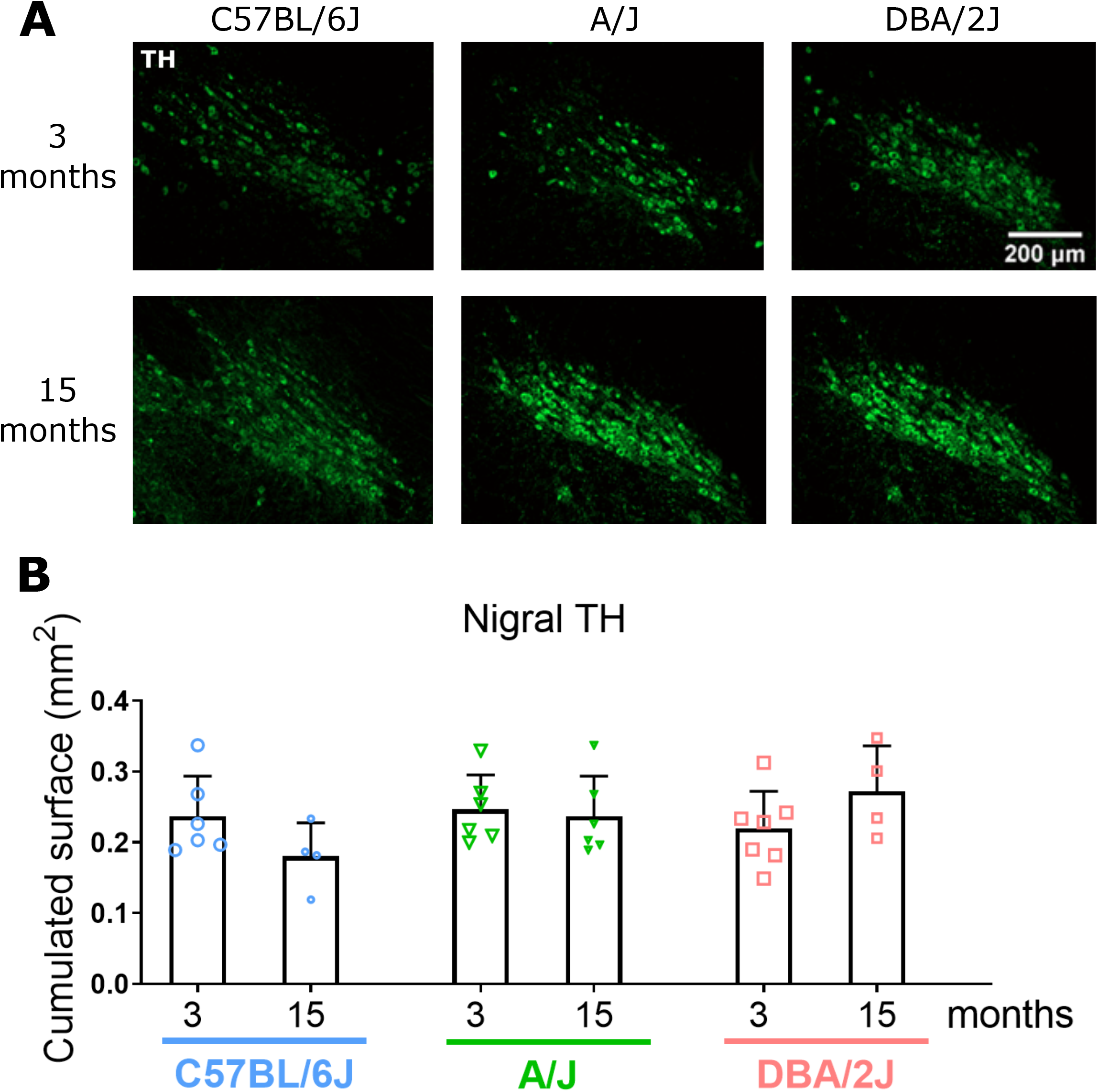
Tyrosine hydroxylase (TH)-positive nigral neurons measured by immunofluorescence in 3- and 15-month-old C57BL/6J, A/J and DBA/2J mice. (A) Images of TH staining, magnification 10X. (B) Quantification of cumulated surface of TH-positive neurons. Data are shown as means ± standard deviation, and analyzed with 2-way ANOVA, followed by Tukey’s multiple comparisons test.

## 4. Discussion

Genetic factors as well as aging modulate an organism’s tissues normal evolution and susceptibility to disease. This is particularly true for the CNS, for which aging is typically associated with functional and, in some instances, structural decline, as well as increased propensity to develop neurodegeneration. The brain is an organ typically impossible to access for biopsies, and thus, for obvious ethical reasons, longitudinal studies are difficult in humans. Thus, mammalian model organisms will continue to play a pivotal role in understanding physiological aging and its effect on disease (Mitchell et al., 2015). The mouse is particularly well suited for this purpose because of the availability of strains with different genetic backgrounds, and the possibility to longitudinally analyze aging study cohorts. Despite this advantage, most studies on the CNS use only one mouse strain (usually a C57BL6 substrain), and few use aging cohorts.

Earlier studies have analyzed CNS features in aging mice, even though many of them only qualitatively (Ingram and Jucker, 1999; Jucker and Ingram, 1997). Lipofuscin accumulation appears to be a consistent feature of the aging mouse brain (Jucker and Ingram, 1997). Neuronal atrophy and reduction, in various brain regions, of dendritic or axonal arborizations have been described in various aged C57BL/6 substrains, but, no loss of neurons in different brain regions (Ingram and Jucker, 1999; Jucker and Ingram, 1997). Age-dependent increase in glial reactivity, transcriptome and DNA methylation changes have also been reported for mouse brain (Jucker and Ingram, 1997). In a recent study (Shaerzadeh et al., 2020), the number of TH-positive neurons in the SN of C57BL/6 mice (substrain not specified) did not change after 1 months of age up until 24 months. Interestingly though, the number of microglia surrounding these neurons and their contacts with them increased with age, possibly reflecting an increased trophic support of microglia for these neurons as they age. Finally, many genetically modified mice have shown age-dependent alterations of various structural, molecular, and functional CNS features (Heng, 2017).

Very little is known about age-related alterations in the nigro-striatal circuit in mice of different genetic backgrounds. The nigro-striatal circuit in the CNS is a key neuronal pathway that connects the dopaminergic neurons of the SN to the dorsal striatum. It is one of the major dopaminergic circuits of the CNS and is evolutionary well conserved (Vogt Weisenhorn et al., 2016). Because of its central role in the control of voluntary motor movement and its relevance to PD (Raza et al., 2019; Vogt Weisenhorn et al., 2016), it is at the center of neuroscience research. Hence, in this study, we assess the integrity of the nigro-striatal circuit in aging cohorts of three common laboratory inbred mouse strains, C57BL/6J, A/J, and DBA/2J. We picked these three strains, not only as they are commonly used, but also because there is substantial information already available on their behavior, in particular motor behavior (Brooks et al., 2004; Ingram and Jucker, 1999; Ingram et al., 1981; Kollmus et al., 2020), and their susceptibility to PD neurotoxins (Vogt Weisenhorn et al., 2016; Yin et al., 2011). However, no comparative analysis has been done yet of the effect of both strain and age on the biochemical and structural properties of the nigro-striatal circuit. In addition, these 3 strains are, either in pair-wise combination or in combination with other strains, founders for a number of mouse genetic reference populations, such as the BXD mice (Ashbrook, 2019; Peirce et al., 2004), the C57BL/6J – A/J Chromosome Substitution mice (Nadeau et al., 2012), and the Collaborative Cross mice (Threadgill et al., 2011). Thus, we aim our study to be a stepping stone for further ones aimed at deeper analysis of nigro-striatal circuit properties, such as susceptibility to PD toxins across strains and age, and mapping of genetic regulators of nigro-striatal response to aging and stressors.

We analyzed the striatal dopamine levels and the morphological integrity of the nigro-striatal circuit. We thus set a baseline of biochemical and morphological nigro-striatal features in C57BL/6J, A/J, and DAB/2J mice at 3, 9, and 15 months of age. We decided to focus on these features as behavior studies are already widely available (see references above), and because biochemical and morphological features are ultimately the substrates of CNS functionality. A large part of our study looks at morphological features by immunohistochemical techniques. Quantitative image analysis of immunohistochemically labeled tissue sections is widely used in neuroscience and other fields to measure protein expression and morphological density of immunopositive structures (Crowe and Yue, 2019). Standardization of tissue preparation (extraction, fixation, and work-up), staining protocol, and “blinded” image acquisition enable trustworthy analyses of cellular structures and proteins at a detailed anatomical scale (Guirado et al., 2018; Jensen, 2013).

Using these approaches, we made the key observation that all dopaminergic markers analyzed in the striatum showed, in each mouse strain, a different evolution across age. The number of TH-positive neurons in the SN though remained constant up to 15 months. Our observations show that a great degree of plasticity exists at the level of the axonal projections of these neurons, even at older age. We had only a limited number of mice for the 15-month-old DBA/2J mice-however, the observation that all markers we measured were increased in this group of mice, and not just one, makes us feel confident about the reliability of these results. The observation that some features of striatal axons of SN dopaminergic neurons are counter-intuitively increasing with age in some strains (A/J, DBA/2J) may reflect processes that are aimed at compensating for aging. Because these axonal structures are the first and foremost ones affected in PD (Kordower et al., 2013; O’Keeffe and Sullivan, 2018), molecular dissection of the underlying genetic mechanisms may open up new insights that have important implications for the understanding of the disease.

Few studies look at nigro-striatal circuit features in different strains of mice or other species. Some of them (Vadasz et al., 1987; Zaborszky and Vadasz, 2001) have demonstrated variable mesencephalic TH enzyme activity in different mouse strains that, curiously, seemed to corrrelate inversely with actual TH-positive neuron number. A proteomics profiling study showed wide ranging differences in striatal proteins between C57BL/6J and DBA/2J mice (Parks et al., 2019). In one of our recent studies we have shown, using a trait mapping approach with a collection of Collaborative Cross mice, that the subunit a6 of collagen 4 acts as a probable novel regulator of striatal dopamine level and axonal branching of dopaminergic neurons (Thomas, 2020).

Studies that analyze the effect of age on nigro-striatal circuit’s features in mice are also not very numerous. Age was reported to affect different features of dopamine release in C57Bl6/J mice (Arvidsson et al., 2014), even though it does not affect total dopamine level and morphological features of the striatum, as shown in the present study. Recently, by comparing the ventral midbrain gene expression profile in the same 3 mouse strains as in this study, we have shown that the transcription factor Pituitary Tumor Transforming Factor Gene (Securin) orchestrates expression profiles in this brain region over age (Gui, 2020).

Several studies indicate that age-dependent evolution of the nigro-striatal circuit varies between rat strains. While in Wistar rats, striatal dopamine levels and TH activity diminish with age (De La Cruz et al., 1996; Watanabe, 1987), Fischer 344 (F344) rats show an age-dependent increase of dopaminergic striatal innervation (Ishida et al., 2007). However, these rats show age-related decline in DA uptake (Hebert and Gerhardt, 1999), which is likely due to a decrease in DAT located at the plasma membrane rather than a decrease of total DAT (Salvatore et al., 2003). In contrast, in aged outbred Sprague-Dawley rats, the overall level of DAT decreases with age, while a more detailed look indicates an age-dependent complex resdistribution of post-translationally modified forms of DAT between subcellular compartments (Cruz-Muros et al., 2009). Also in sprague Dawley rats, VMAT2 shows an age-dependent subcellular redistribution of its different post-translationally modified forms, whereas the total level of VMAT2 doesn’t change (Cruz-Muros et al., 2008). In rhesus monkeys, no age related loss of TH neurons in the SN was observed, but an age-related decline of striatal dopamine and TH fiber density, as well as a reduced capacity to compensate for MPTP-induced lesions (Collier et al., 2007).

Taken together, all these observations show that genetic background as well as age modulate various structural and functional properties of the nigro-striatal circuit in rodents. They also show that this circuit presents with a remarkable plasticity, which may allow it to respond to age-related stress and exposure to environmental toxins. Several studies support this assertion. DBA/2J mice are more resistant to the neurotoxin MPTP than C57BL/6J mice (Hamre et al., 1999). Strains from the genetic BXD reference population, which are derived from C57BL/6J and DAB/2J founders, show a scale of responses to the toxin paraquat (Torres-Rojas et al., 2020). The nigro-striatal circuit of 2-to 3-month-old C57BL/6J mice is more resilient and shows better recovery after MPTP than that of 12-month-old mice of the same strain (Date et al., 1990; Filipov et al., 2009).

Several lines of evidence indicate that, with aging, the functionality and integrity of nigral projections into the striatum also decrease in humans (Karrer et al., 2017; Volkow et al., 1996a; Volkow et al., 1996b). These striatal axonal projections are among the very first CNS structures to decay in PD, and they do so before classical neurological symptoms are observed (Kordower et al., 2013; O’Keeffe and Sullivan, 2018). It is tempting to speculate that individual variability in the different levels of functionally important striatal proteins, such as those measured in mice the present study, may modulate susceptibility to age- and PD-related challenges. Again, it is mouse models that provide evidence that this is indeed the case. Genetically altered C57BL/6 mice (substrain not specified) with reduced levels of VMAT2 show age-dependent decline of dopaminergic axon loss, and enhanced susceptibility to MPTP, whereas mice of that strain with overexpression of VMAT2 are protected against MPTP-induced striatal damage (Lohr et al., 2014). Similarly, mice (backcrossed 5 generations on C57BL/6J) with reduced DAT expression are partially resistant to paraquat neurotoxicity (Rappold et al., 2011), and mice (mixed C57BL/6J-l29sVj background) completely lacking DAT are resistant to MPTP neurotoxicity (Gainetdinov et al., 1997). Evidence suggest that the absence of DAT is protective by preventing entry of paraquat and of the toxic metabolite MPP+ into neurons (Shimizu et al., 2003; Zimmer et al., 2000). Interestingly, genetic variants in the DAT gene sequence modulate PD risk after exposure to pesticides, and this effect is gene dosage dependent (Kelada et al., 2005; Ritz et al., 2016). The effect of these genetic variants on DAT level and function remains to be determined. Two-to 3-month-old C57BL/6 mice (substrain not specified) lacking the *Aldh1a1* gene showed alterations in dopamine metabolism, reuptake, and DAT functionality (Anderson et al., 2011). *Aldh1a1*-expressing SN neuron subpopulations in mice (strain not specified) with local overexpression of *Aldh1a1* are protected from α-synuclein toxicity, whereas the same neurons lacking Aldh1a1 expression are more susceptible to it (Liu et al., 2014). Interestingly, in PD, the level of ALDH1A1 are greatly reduced in the nigro-striatal circuit, which is believed to lead to accumulation of toxic dopamine metabolites (Grunblatt and Riederer, 2016).

Thus, all this evidence shows that mice and other rodents will remain essential tools in biomedical research, in particular neuroscience (Green et al., 2011). Mouse model use in translational neuroscience has been questioned (Manger et al., 2008), in particular because of their reported lack of predictive power for what actually happens in the clinic (King, 2018; Potashkin et al., 2010). In PD though, they have been essential in supporting the development of dopamine replacement therapies (Duty and Jenner, 2011), which is currently one of the main therapeutic options for this disease. Several thoughtful suggestions have been put forward to improve the usefulness of mouse models in translational neuroscience, and one of them is to enlarge the number of background strains in mouse studies (2009; Perrin, 2014). Indeed, the lack of predictive power may not come as a surprise since most neuroscience studies are done on just one or a limited number of strains of mice, and mostly in cohorts of a particular age which can be considered “young adults” (usually around 3-month-old). On the one hand, large mouse modeling studies that incorporate several different strains and ages are daunting and bound to be very resource-and time intensive. On the other hand, the huge costs associated with developing experimental therapies that ultimately fail, and the urgency to address some of the most devastating neurological diseases of our time, such as PD, should encourage such studies to be designed. In this context, our study provides a first detailed look into the genetic background- and age-associated variabilities of features in the nigro-striatal circuit, a structure that is most affected in PD. Thus, it underscores the importance of considering specific strains, or combination of strains, as well as age of mice in designs of studies around this neuronal circuit in health and disease.

## Ethics statement

The protocol for mice bred at University of Luxembourg was reviewed and approved by the Animal Experimentation Ethics Committee (AEEC). For the mice bred in Helmholtz Centre for Infection Research (Braunschweig, Germany), the protocol was reviewed and approved by the ‘Niedersächsisches Landesamt für Verbraucherschutz und Lebensmittelsicherheit, Oldenburg, Germany’ (Permit Number: 33.19-42502-05-19A394).

## Author contributions

MB, PG and LS conceived the project with input from AM, KS, RB and MM. MT, MK, BPR, YG, MB and PG designed the experiments and analysis. MT, MK, PG, and MB prepared mouse tissues. MT, MK, and BPR performed immunofluorescence experiments. CJ performed GC-MS experiment. DC, HK and KS supported mouse breeding. MT, MK, BPR, YG, CJ, MM, LS, PG and MB analyzed the results. MT and MB wrote the manuscript. All authors read and approved the final manuscript.

## Acknowledgements

Acknowledgements: LS and MB thank the Luxembourg National Research Fund (FNR) for funding support (FNR CORE C15/BM/10406131 grant). MM thanks the FNR for funding support (FNR PEARL P16/BM/11192868 grant). KS thanks the support by intra-mural grants from the Helmholtz-Association (Program Infection and Immunity) and the animal caretakers at the Central Animal Facilities of the HZI for maintaining the mice.

## Abbreviations

ALDH1A1: aldehyde dehydrogenase 1 family member A1
BSA: bovine serum albumin
CNS: central nervous system
DAT: dopamine transporter
GC-MS: gas chromatography - mass spectrometry
MPTP: 1-methyl-4-phenyl-1,2,3,6-tetrahydropyridine
PBS: phosphate-buffered saline
PD: Parkinson’s disease
PFA: paraformaldehyde
SN: substantia nigra pars compacta
TH: tyrosine hydroxylase

