## Supplementary Fig.S1 for "Strain- and age-dependent features of the nigro-striatal circuit in three common laboratory mouse strains, C57BL/6J, A/J, and DBA/2J - *Implications for Parkinson’s disease modeling*"

### Supplementary Figures

Supplementary Figure S1.

Striatal Aldehyde Dehydrogenase 1 Family Member A1 (ALDH1A1) measured by immunofluorescence in mice. (A) Tile acquisition of images of ALDH1A1 and tyrosine hydroxylase co-staining in the striatum of a 3-months-old C57BL6/J mouse, magnification 10X. (B) Tile acquisition of images of ALDH1A1 staining in the striata of 3-, 9- or 15-months-old C57BL/6J, A/J or DBA/2J mice, magnification 10X.
