## Supplementary figures and images for "Strain- and age-dependent features of the nigro-striatal circuit in three common laboratory mouse strains, C57BL/6J, A/J, and DBA/2J - *Implications for Parkinson’s disease modeling*"

### Supplementary Fig.S1

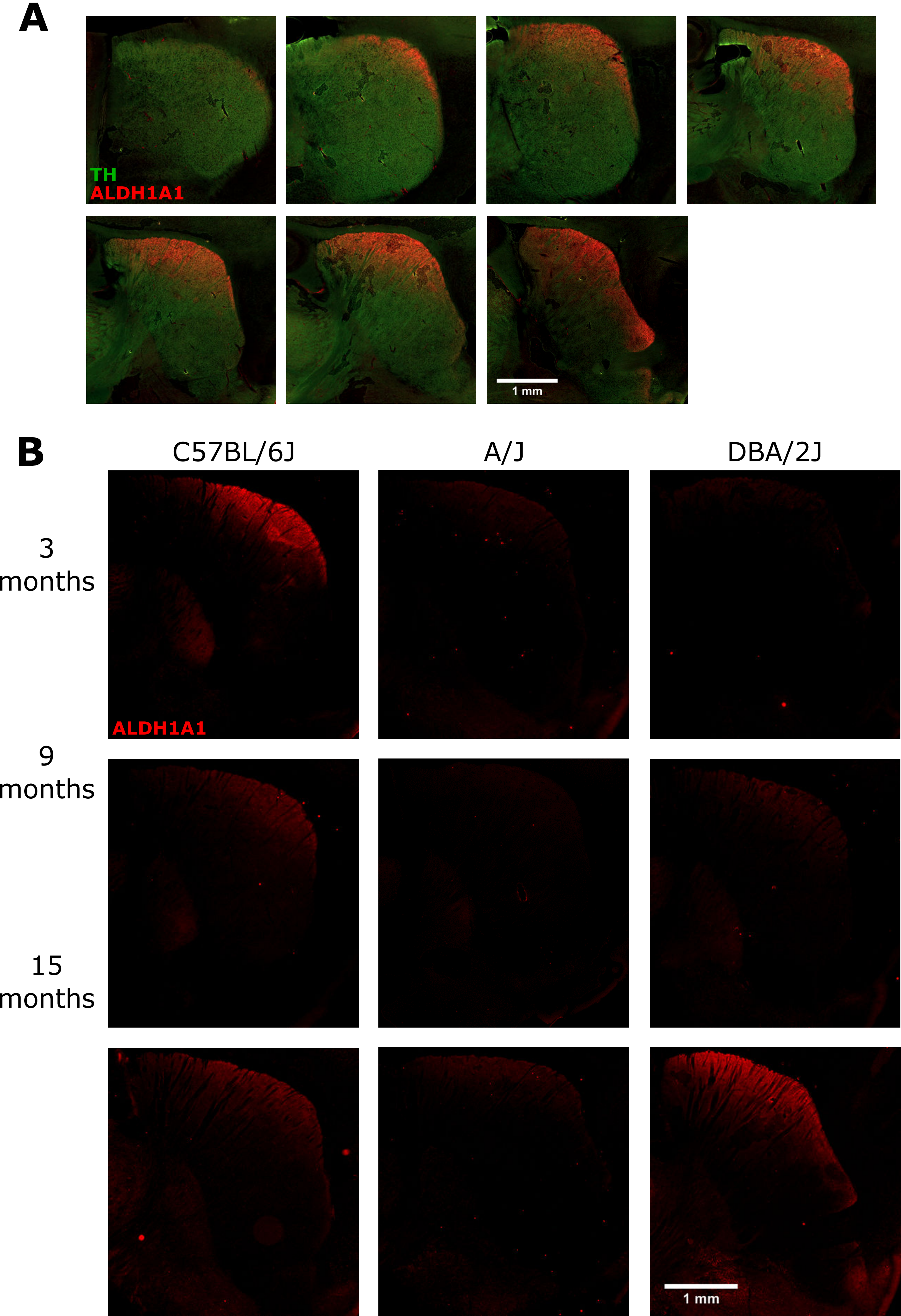
